# No evidence for trans-generational immune priming in *Drosophila melanogaster*

**DOI:** 10.1101/2023.04.25.538340

**Authors:** R. Radhika, Brian P. Lazzaro

## Abstract

Most organisms are under constant and repeated exposure to pathogens, leading to perpetual natural selection for more effective ways to fight-off infections. This could include the evolution of memory-based immunity to increase protection from repeatedly-encountered pathogens both within and across generations. There is mixed evidence for intra- and trans-generational priming in non-vertebrates, which lack the antibody-mediated acquired immunity characteristic of vertebrates. In this work, we tested for trans-generational immune priming in adult offspring of the fruit fly, *Drosophila melanogaster*, after maternal challenge with 10 different bacterial pathogens. We focused on natural opportunistic pathogens of *Drosophila* spanning a range of virulence from 10% to 100% host mortality. We infected mothers via septic injury and tested for enhanced resistance to infection in their adult offspring, measured as the ability to suppress bacterial proliferation and survive infection. We categorized the mothers into four classes for each bacterium tested: those that survived infection, those that succumbed to infection, sterile-injury controls, and uninjured controls. We found no evidence for trans-generational priming by any class of mother in response to any of the bacteria.

## INTRODUCTION

Most organisms are under constant and repeated exposure to pathogens and parasites. Since offspring are likely to share an environment with their parents and therefore experience immune challenges similar to those their parents did, natural selection may favor the evolution of mechanisms that allow immune experiences to be transferred across generations. Trans-generational immune priming (TGIP) is a phenomenon where parental exposure to a parasite or pathogen results in enhanced defense against that parasite in their offspring (Roth et al. 2018; Gourbal et al. 2018). In vertebrates, this can be achieved with maternal transfer of antibodies (Hasselquist and Nilsson 2009) but because invertebrates lack antibody-mediated immunity, potential mechanisms for TGIP in invertebrates would need to rely on altered gene expression or tissue development. There has been some evidence for TGIP in insects (reviewed in Gourbal et al. 2018) but it is unclear how widespread the phenomenon is, and most insects in which TGIP has been observed are not particularly tractable for experiments that would elucidate the mechanistic basis. In the present work, we test for TGIP in *Drosophila melanogaster*, which is highly amenable to mechanistic study, but find no evidence of elevated immunity in adult offspring after maternal infection with ten different bacterial pathogens. Given that the fruit fly is a model system widely used for many fields of research, including immunology-related research, we believe it would be quite important to know if this phenomenon is exhibited b the species.

TGIP was first described in vertebrates when it was discovered that mothers pass to their offspring immune memories of pathogens to which they have been exposed in the form of highly specific antibodies via milk, blood or maternal deposition in eggs (Boulinier and Staszewski 2008; D. Hasselquist, J.-Å. Nilsson 2009). If the offspring became subsequently infected with a pathogen against which they had been maternally primed, they induced a stronger immune response than they would have if they had been unprimed. TGIP has been reported in some insect taxa such as bumble bees (Sadd et al. 2005), mealworm beetles (Moret 2006), flour beetles (Roth et al. 2009), meal moths (Tidbury, Pedersen, and Boots 2011), and cockroaches (Faulhaber and Karp 1992), as well as a few other invertebrates such as *Daphnia* (Little et al. 2003), a copepod *Macrocyclops albidus* (Kurtz and Franz 2003) and *Biomphalaria glabrata* snails (Portela et al. 2013). In comparison to vertebrates, TGIP among invertebrates has been shown to impart offspring with a more ‘generalist’ priming that is not specific to the pathogen their parents were exposed to, possibly due generic upregulation of immune systems as opposed to specific resistance to the maternal infection (Dhinaut et al 2017; Gourbal et al. 2018). Among insects, TGIP seems to be more common in orders like Coleoptera, Lepidoptera and Hymenoptera than Diptera (summarized in Pigeault et al. 2016, Tetreau et al. 2019). It remains uncertain precisely how widespread TGIP is among insects and other invertebrate animals, and the mechanistic basis for TGIP is largely unknown (but see Tetreau et al. 2019).

A previous study in *Drosophila melanogaster* indicated that material infection with two different bacteria did not increase the ability of adult offspring to survive infection with those same two bacteria (Linder and Promislow 2009). In this study, we expand that work, testing for TGIP in *D. melanogaster* after maternal infection with ten different bacterial pathogens that span a range of virulence (Table 1) and measuring both survivorship after infection and capacity to suppress pathogen proliferation in the offspring. In order to test whether the mother’s infection outcome was predictive of her ability to prime offspring, we categorized the mothers based on whether they survived their infection or died from it and contrasted resistance in the adult offspring of mothers from both groups to resistance of offspring of mothers that received either a sterile injury or were uninjured. We also tested whether there was a temporal dynamic to maternal priming, evaluating resistance in offspring produced within 24 hours of maternal infection, 2-3 days after maternal infection, or 4-5 days after maternal infection. We found no difference in the ability of offspring to survive infection or suppress pathogen growth after maternal infection with any of the ten bacteria tested in any of the data subsets, nor when all of the data are combined, so we conclude that *D. melanogaster* mothers are unable to trans-generationally prime their adult offspring against bacterial infection.

**Table 1:** List of bacteria used to infect *D. melanogaster* in our experiment

| Name of bacteria | Gram type | Virulence | Reference |
| --- | --- | --- | --- |
| *Providencia sneebia* | Gram negative | high | Juneja *et al.*2009 |
| *Pseudomonas aeruginosa* | Gram negative | high | Stover *et al.*2000 |
| *Providencia rettgeri* | Gram negative | moderate | Galac *et al.*2011 |
| *Escherichia coli* | Gram negative | low | Troha and Im *et al.*2018 |
| *Serratia marcescens* | Gram negative | low | Troha and Im *et al.*2018 |
| *Staphylococcus aureus* | Gram positive | high | Troha and Im *et al.*2018 |
| *Enterococcus faecalis* | Gram positive | moderate | Troha and Im *et al.*2018 |
| *Bacillus subtilis* strain 3610 | Gram positive | moderate | Cai *et al.* 2017 |
| *Bacillus subtilis* strain 168 | Gram positive | moderate | Cai *et al.*2017 |
| *Micrococcus luteus* | Gram positive | low | Troha and Im *et al.*2018 |

## METHODS

### Fruit Fly Husbandry and Infection Protocol

*D. melanogaster* from the wild-type laboratory strain, Canton S, were used for all of our experiments. Flies were reared in mixed-sex populations to allow mating at a density of 100 eggs per vial (±10%) under a 12:12 hour light:dark cycle at 25°C and 50-60% relative humidity. On the 12th day after egg collection (when flies are 2-3 days old as adults), female flies were given one of the following three treatments: bacterial infection, sterile injury, or uninjured control. Infections were carried out by exposing flies to mild CO_2_ anaesthesia and pricking on the lateral side of their thorax with a thin needle (0.1 mm) dipped in a bacterial suspension (Khalil et al 2015). To prepare bacterial suspensions, bacteria were cultured overnight in Luria-Bertani broth (LB) at 37°C to form a primary culture. The primary culture was used to seed a secondary culture in LB that was again grown at 37°C for ∼3 hrs. Bacteria from secondary culture were pelleted by centrifugation at 7000 RPM for 5 min and re-suspended in phosphate-buffered saline (PBS) to achieve an A_600_ of 1.0. This slurry was used for infections. For each of the bacterial strains we used, this method delivered a few thousand bacterial cells into the septic wound. Injury controls were pricked with a needle dipped in PBS while uninjured controls were merely exposed to CO_2_ anesthesia. We tested infection with ten species/strains of bacteria that vary in virulence from 10% to 100% host mortality over the experimental period (Table 1). Many of these bacteria were originally isolated as natural infections of *D. melanogaster* and all are commonly used in *Drosophila* infection research.

Flies from each infection treatment were housed individually in vials for oviposition and eggs were collected to subsequently test for priming of the offspring. Eggs were collected over three time intervals: 0-24 hours after maternal infection, 2-3 days after maternal infection, and 4-5 days after maternal infection. For each of these three sets, eggs collected from infected mothers were segregated into 2 groups: those that were laid by mothers that succumbed to infection within the duration of oviposition and those that were laid by mothers that survived infection until the end of oviposition period. Thus, four sets of eggs could be obtained for each window of oviposition for each bacterium tested: from mothers who were uninjured, from mothers who received a sterile injury, from mothers who received infection and survived, and from mothers who received infection and died. Most bacteria used for experiments did not kill all flies by the end of Day 5. Hence, we were able to obtain eggs from all three windows of oviposition. For bacteria that killed all mothers before the end of a given egg collection period, fewer oviposition windows were sampled. In order to test for TGIP, progeny that developed from the collected eggs were either infected with the same bacteria used to infect their mothers or were injured with a sterile needle. Pathogen burden and post-infection survivorship were compared between the offspring of infected mothers versus offspring from mothers who were either sterilely wounded or anesthetized only.

### Survivorship and Bacterial count assays

Resistance to infection in the offspring was tested on the 12th day after egg collection. This is approximately 3 days after offspring eclosed from pupal case as adults. For survivorship assays, 50 flies were infected with each bacterium as described above and 30 were pricked with a sterile needle to generate injury controls. Mortality rates were observed for five days after infection. Our injury controls suffered no mortality during the window of observation. For bacterial count assays, 15-20 flies from each treatment were individually pulverized in 500 μl of PBS using a steel ball and the homogenate was plated on LB agar using a robotic spiral plater (Don Whitley Scientific) at 0, 4, 12, 24 and 48 hours post-infection. Plates were incubated overnight at 37°C and the bacterial colonies that grew were counted using the ProtoCOL automated counter (Microbiology International). The natural flora of *D. melanogaster* reared under lab conditions grows very slowly on LB at 37°C and does not yield visible colonies on plates during the experimental period. The bacteria used for experimental infections were confirmed by colony color and morphology on the plates derived from infected flies. Control homogenates from sterilely wounded flies did not yield visible colonies. Each experiment was performed thrice to generate three biologically independent blocks of data.

### Data analysis

We evaluated survivorship data with a Kaplan-Meier analysis using the package ‘*survminer’* implemented in R. Since we had three independent blocks for each experiments, we first tested for among-block variation in each of the treatments across all three blocks using both Kaplan-Meier and Cox Proportional Hazards analyses. In no case did we find statistically significant variation among blocks within our treatments so we combined all blocks for a final survivorship analysis using Kaplan Meier, and these are the results that are presented.

Bacterial count data were analyzed using analysis of variance (ANOVA) implemented in JMP. A mixed effects model was built with block as a random factor and with time post-infection and maternal condition as fixed factors. Post-infection time had five levels (0, 4, 12, 24 and 48 hours) and maternal condition had four levels (infected-dead, infected-alive, sham infected, uninjured control).

### Bacteria used for experiments

To survey TGIP in these experiments, we exclusively used pathogens that are natural opportunistic pathogens of the fruit fly. We were also keen to see if the amount of virulence the bacteria had against our flies influenced their ability to prime offspring. We reasoned that while infections with bacteria that cause higher virulence might provide a greater incentive for mothers to prime their offspring, such situations leave mothers with fewer resources to invest in reproduction related mechanisms, a common trade-off observed in many insect and non-insect taxa (Schwenke et. al. 2015, French et. al. 2007; Brokordt et. al. 2019; Xu et. al. 2012), thus impairing their ability for TGIP. In order to account for both of these possibilities, we selected ten species/strains of bacteria to span a range of virulence against the host (Table 1).

## RESULTS

We tested for trans-generational immune priming in *Drosophila melanogaster* by evaluating whether progeny of infected mothers and were more resistant to bacterial infection than progeny of uninfected mothers, where resistance was defined as the ability to survive infection and suppress pathogen proliferation. We evaluated potential trans-generational priming after maternal infection with 10 different bacteria that cover a full range virulence from nearly benign to completely lethal (Table 1). In addition to evaluating the full data set, we tested for a potential temporal component of maternal priming by binning offspring produced withing 24 hours, 2-3 days, or 4-5 days after maternal infection. To additionally test whether maternal infection outcome predicted offspring priming, we separately evaluated the offspring of mothers who died from their infections during the oviposition window versus offspring of surviving mothers.

The offspring of mothers from all four categories (infected-died, infected-survived, sterile injury, uninjured) had indistinguishable survival profiles in all three of the oviposition windows (0-24 hours, 2-3 days, or 4-5 days post-infection) for each of the 10 bacteria tested (Figure 1; Kaplan-Meier p >> 0.05 in all cases). There was also no hint of any difference in survivorship when data were pooled across oviposition time windows or when offspring from mothers who died from or survived infection were combined (data not shown; Kaplan-Meier p >> 0.05 in all cases). Thus, we found no evidence that infected mothers prime their offspring in a way that results in increased probability that adult offspring will survive the same infection, irrespective of the mother’s infection outcome or the duration of her infection prior to oviposition.

**Figure 1:**
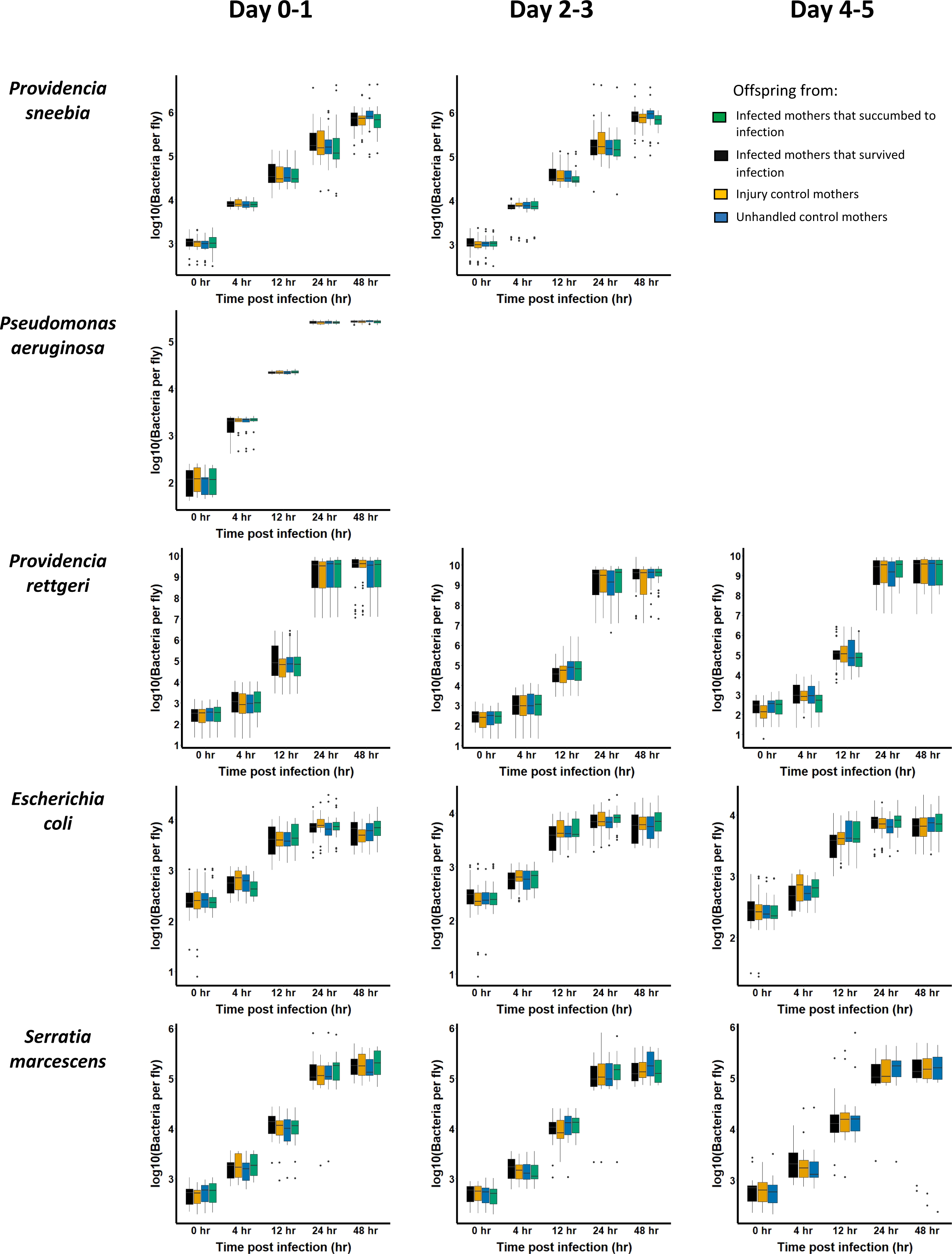
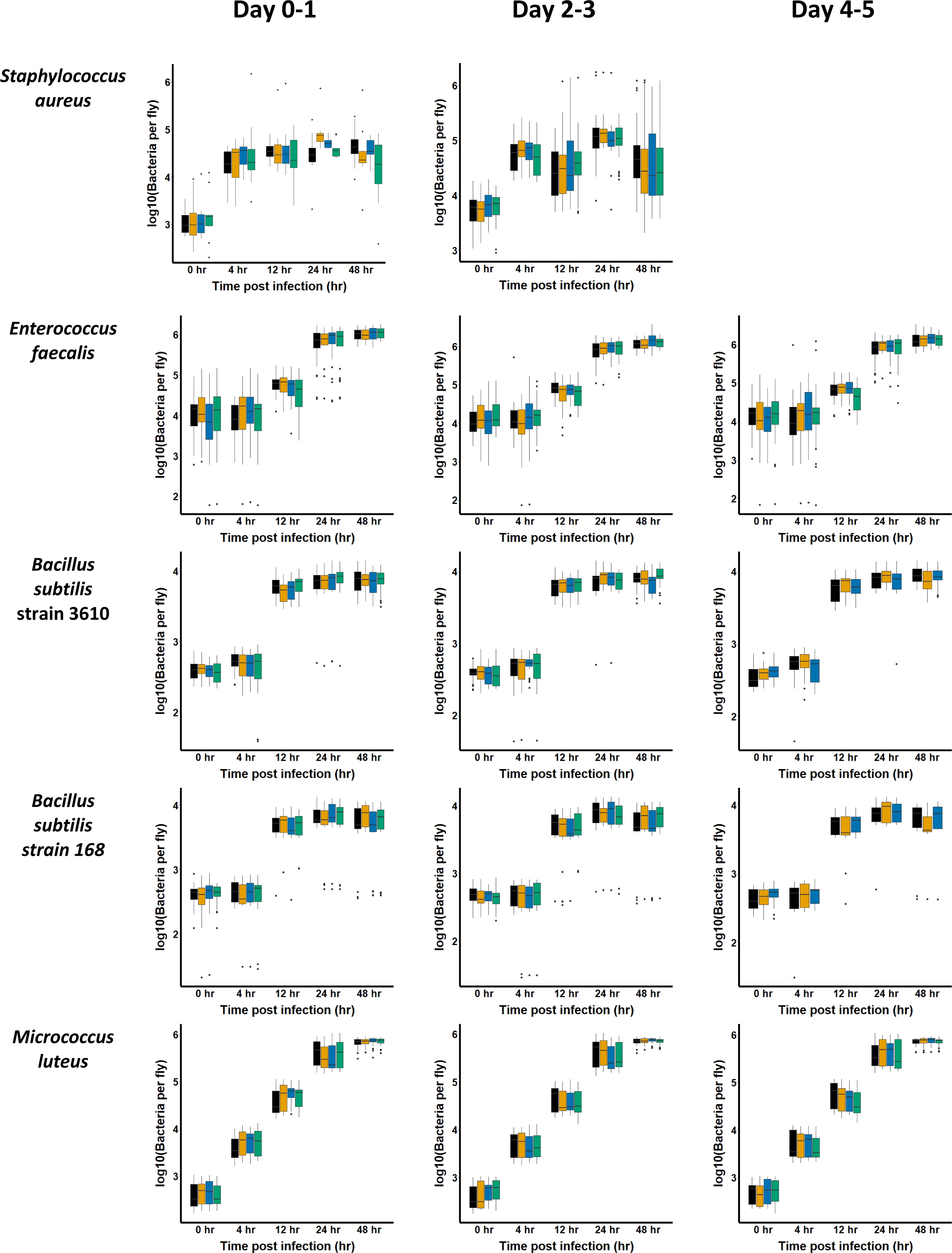
Survival assays to test for priming among offspring. Survival assays were performed against ten different strains of bacteria. To test for priming against each bacterium, from each of the four maternal treatments (infected and succumbed -green, infected and survived - black, sham infected - ochre, uninjured controls - blue) offspring were infected and their survival rates monitored for several days post infection. Survivorship analysis was performed using *Survminer* package in R.

In order to test whether priming occurred at the level of restricting pathogen proliferation, we measured pathogen burden in the infected offspring from each class of mother at 0, 4, 12, 24 and 48 hours post-infection. As expected, time post-infection was a highly significant predictor of bacterial load after infection with each of the 10 bacteria due to bacterial growth over the course of infection (Figure 2, Table 2; ANOVA p < 0.005 in all cases). However, neither maternal condition nor the interaction between maternal condition and time post-infection were statistically significant predictors of pathogen burden in offspring from any oviposition window after infection with nine of the ten bacteria (Figure 2, Table 2; ANOVA p >> 0.05 in all cases). The sole exception was *Bacillus subtilis* strain 3610, for which the interaction between time post-infection and maternal condition was weakly nominally significant (Table 2, p = 0.034). We also saw no effect of maternal condition or the interaction between maternal condition and time post-infection on offspring pathogen burden when the data were from all oviposition windows were combined, or when offspring from mothers who survived infection or died from infection were pooled (not shown; ANOVA p >> 0.05 in all cases). Thus, we found no compelling evidence that infected mothers prime their offspring in a way that results in increased ability to restrict pathogen proliferation. Considering the survivorship and pathogen burden data in combination, we find no evidence that *D. melanogaster* mothers infected with bacteria prime their offspring for enhanced adult resistance to bacterial infection.

**Figure 2:**
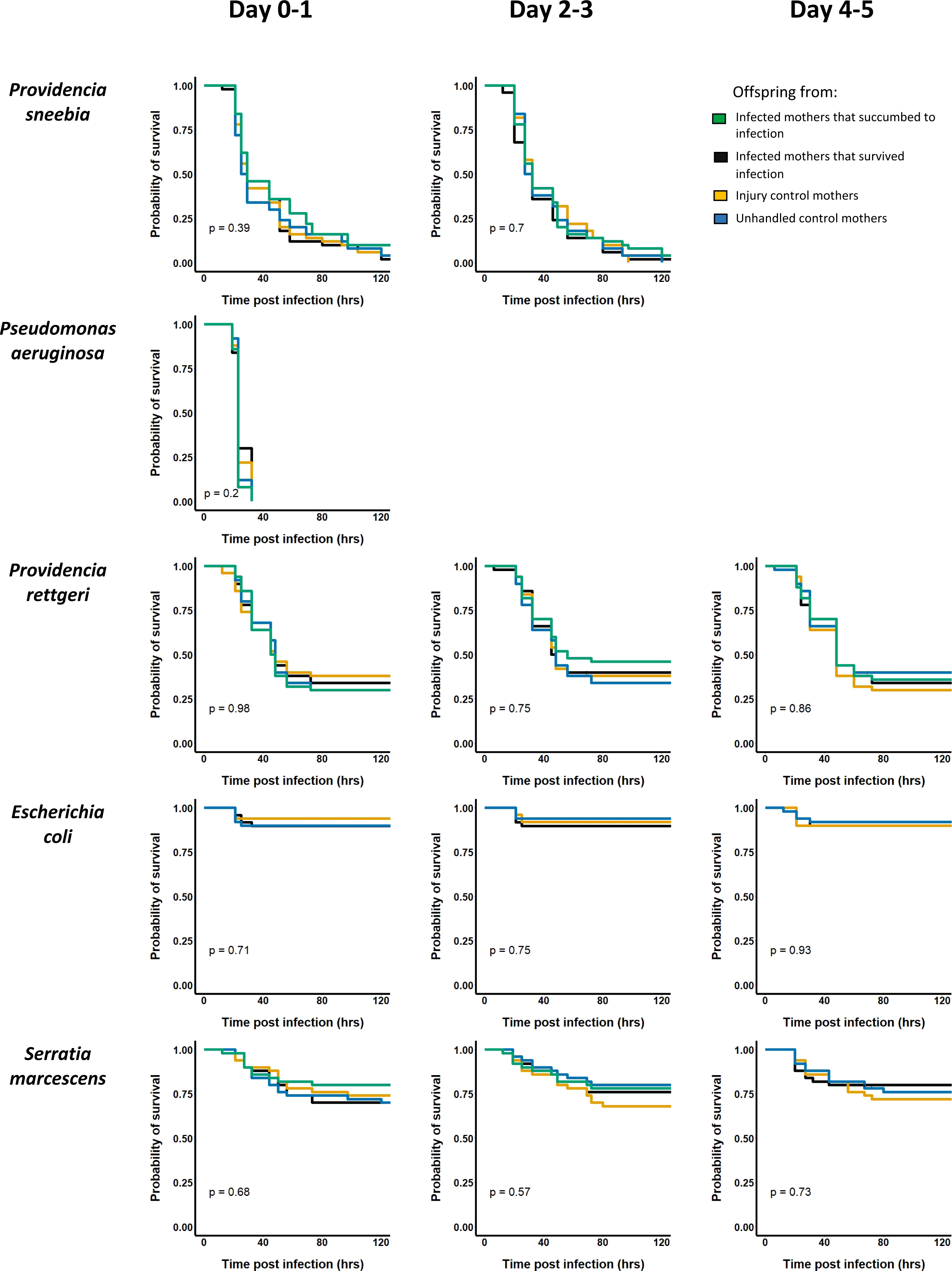
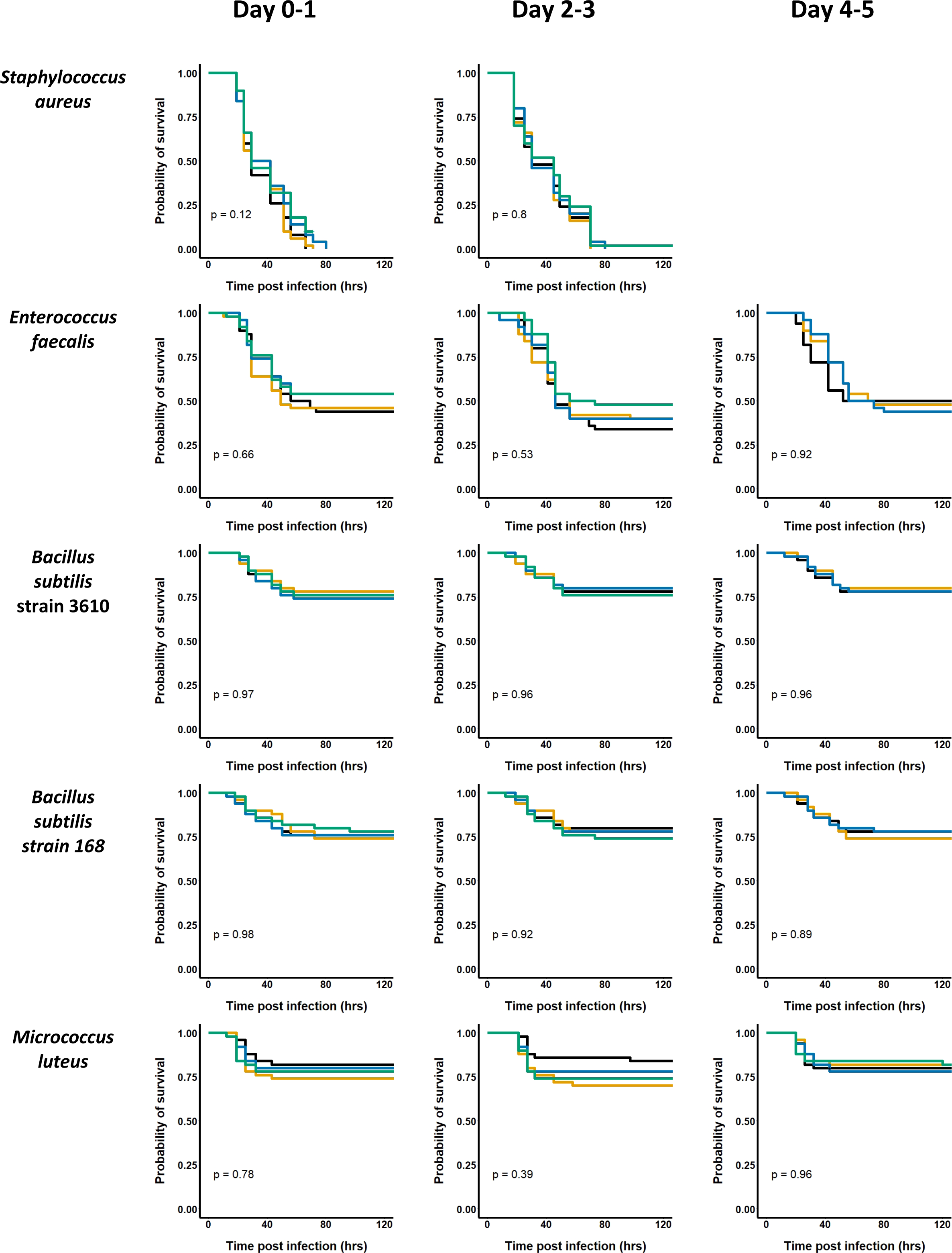
Pathogen load assays to test for priming among offspring. Pathogen load assays were performed after infection with ten different bacteria. For each bacterium, progeny of mothers from each of the four treatments (infected and succumbed – green, infected and survived - black, sham infected - ochre, uninjured controls - blue) were individually crushed in PBS and plated at 0, 4, 12, 24 and 48 hours post infection. Bacterial colonies on plates were counted using an automated algorithm. All experiments per performed thrice to generate three biologically independent blocks of data for each bacterium tested. Statistical analysis of these data is presented in Table 2; maternal condition was not a significant predictor of pathogen burden in offspring in any case.

## DISCUSSION

Despite testing 10 different bacterial strains that vary in virulence to the *D. melanogaster* host, we found no evidence that infected *D. melanogaster* mothers immunologically prime their offspring in a manner that protects the next generation from infection as adults. We evaluated maternal infection with bacteria that were chosen because they are natural pathogens of *Drosophila* or are commonly used in *Drosophila* laboratory experiments. These included both Gram-positive and Gram-negative bacteria that cause a wide range of host mortality (15-100%). We additionally tested for a potential temporal component to priming, separately evaluating offspring produced 0-1, 2-3, and 4-5 days after the maternal infection, but we found no evidence of protective maternal priming at any time post-infection. Finally, we tested for differences in priming ability between mothers that survived infections versus those that succumbed to infections and again found no evidence of protective priming in either group.

There are a handful of caveats to this study as we have performed it. First, because we conducted the experiments with only a single host genotype, it remains possible that other host genotypes might impart some degree of trans-generational priming. However, we note that trans-generational priming was also not observed in an independent study that evaluated bacterial infection in three strains of *D. melanogaster* infected with two species of bacteria, all of which were distinct from those used in our study (Linder and Promislow 2009). Thus, it seems safe to conclude that trans-generational priming is not a trait generally exhibited by *D. melanogaster*. Some of the bacteria used in our study cause very high or very low mortality, meaning there may be limited phenotypic space in which priming could be observed. However, we felt it was important to cover a range of virulences in testing for the phenomenon, and our large sample sizes under the various maternal states should give us adequate power to detect meaningful differences in offspring immunity yet we see no hint of any improvement in offspring immunological potential from infected mothers in any category.

An important caveat is that we tested for priming against infection in adult offspring. This leaves open the possibility that there may be transient priming of larval offspring that dissipates prior to completion of metamorphosis into an adult. Such transient priming of larvae may seem logical, especially since larvae have low dispersal capability and therefore are constrained to remain close to the spot in which they were deposited as eggs. However, adult *Drosophila* have high dispersal capability and infected females may lay their eggs quite far from the location in which they acquired the infection. Furthermore, the larval stage is pre-reproductive, and the full fitness benefit of TGIP would be realized only if protection extended to reproductively competent adult offspring that remain in the high-risk environment. Pigeault et al (2016) performed a metanalysis of existing empirical data on immune priming in insects and found that short lifespan and high dispersal, both of which characterize *Drosophila*, are life history traits associated with low levels of maternal immunological priming across insects. This observation is consistent with a theoretical model produced those authors (Pigeault et al 2016), which indicates that trans-generational immune priming is evolutionarily adaptive only when the offspring both have a high probability of sharing environment with the parent and are strongly protected by the parental investment. Our failure to observe TGIP in the present study is therefore consistent with expectations derived from the life history of *D. melanogaster*.

In vertebrates, heritable immunity to particular pathogens is mediated by specific antibodies that are passed from mother to offspring. However, invertebrate immune systems are generalized and lack specificity. Thus, offspring in invertebrate taxa where TGIP has been observed (e.g., Faulhaber and Karp 1992; Roth et al. 2009; Tidbury, Pedersen, and Boots 2011; Pigeault et al 2016) may acquire a non-specific protection against a wider range of pathogens beyond the one initially infecting the mother (e.g. Moret 2006; Shi et al. 2014; Fisher and Hajek 2015; Dhinaut et al 2017; Ben-Ami 2020). Among the invertebrate taxa that do have reported TGIP, the mechanistic basis of how immune priming is transferred from parents to offspring has been studied only in a few instances. Transfer of epigenetic markers has been shown to be a possible mechanism among tobacco hornworms (Gegner et al 2019). Another possible mechanism of priming offspring involves the transfer of microscopic fragments of bacteria from infected mothers to eggs (Freitak et al 2014) along with maternal immune effectors (Dhinaut et al 2017).

TGIP among insects is measured either as increased activity of the immune system (termed Transgenerational Effect on Immunity by Pigeault et al (2016)) or as increased capacity to control and survive infection (termed Transgenerational Effect on Resistance by Pigeault et al 2016). The diversity of insects from which one or both measures of TGIP has been observed might. We emphasize that these are distinct measures that may have different underlying mechanistic bases, and that either one could occur without the other (e.g., Shi, Lin, and Hou 2014; Barribeau, Schmid-Hempel, and Sadd 2016). TGIP in one form or the other has been reported in phylogenetically diverse insects, which could imply that TGIP is widespread. However, selective publication bias could result in over-reporting of examples where TGIP is affirmatively found if such studies are more likely to be published than studies that do not find TGIP, or if studies that fail to detect TGIP are published in venues that have lower visibility. The problem may be exacerbated by investigator bias if experiments to test for TGIP are abandoned when preliminary data show no evidence of an effect. There, thus, is some risk that the scientific literature implies TGIP to be more ubiquitous than it actually is. It remains unclear how common TGIP is among insects and other invertebrates, and whether it has recurrently evolved independently in the taxa in which it has been reported or whether lineages such as *D. melanogaster* that show no evidence of TGIP reflect evolutionarily derived losses. Resolving this will require a systematic study of TGIP in taxa sampled across the invertebrate tree of life, with both positive and negative outcomes reported and mechanistic bases determined where possible.

## ACKNOWLEDGMENTS

We thank all members of the Lazzaro lab for feedback on the project and manuscript.

## FUNDING ACKNOWLEDGMENT

This research was supported by NIH grant R01 AI141385

